# Dispersal limitation and thermodynamic constraints govern spatial structure of permafrost microbial communities

**DOI:** 10.1101/265132

**Authors:** Eric M. Bottos, David W. Kennedy, Elvira B. Romero, Sarah J. Fansler, Joseph M. Brown, Lisa M. Bramer, Rosalie K. Chu, Malak M. Tfaily, Janet K. Jansson, James C. Stegen

## Abstract

Understanding drivers of permafrost microbial community composition is critical for understanding permafrost microbiology and predicting ecosystem responses to thaw, however studies describing ecological controls on these communities are lacking. We hypothesize that permafrost communities are uniquely shaped by constraints imposed by prolonged freezing, and decoupled from factors that influence non-permafrost soil communities. To test this hypothesis, we characterized patterns of environmental variation and microbial community composition in permafrost across an Alaskan boreal forest landscape. We used null modeling to estimate the relative importance of selective and neutral assembly processes on community composition, and identified environmental factors influencing ecological selection through regression and structural equation modeling (SEM). Proportionally, the strongest process influencing community composition was dispersal limitation (0.36), exceeding the influence of homogenous selection (0.21), variable selection (0.16), and homogenizing dispersal (0.05). Fe(II) content was the most important factor explaining variable selection, and was significantly associated with total selection by univariate regression (R^2^=0.14, p=0.003). SEM supported a model in which Fe(II) content mediated influences of the Gibbs free energy of the organic matter pool and organic acid concentration on total selection. These findings reveal that the processes shaping microbial communities in permafrost are distinct from those in non-permafrost soils, as the stability of the permafrost environment imposes dispersal and thermodynamic constraints on permafrost communities. Models of permafrost community composition will need to account for these unique drivers in order to predict community characteristics across permafrost landscapes, and in efforts to understand how pre-thaw conditions will influence post-thaw ecological and biogeochemical processes.

## Introduction

Permafrost is defined as ground that has remained below 0 °C for two or more consecutive years ^1^. Because this definition is solely based on a condition of ‘ground climate’ ^2^, permafrost-affected soils can span a diverse range of soil types and be highly varied in geography, geology, physicochemistry, and microbiology. Indeed, permafrost environments account for approximately 16 % of Earth’s soil environments ^3^, spanning much of the terrestrial Arctic and subarctic ^4^, ice-free areas of Antarctica ^5^, and high-elevation regions in both the northern and southern hemispheres ^4,6^. Collectively, these soils represent an important microbial ecosystem ^7^ and a globally significant pool of sequestered carbon ^3,8^, which is being mobilized as climate warming increases permafrost thaw ^9^. While the fate of this carbon remains uncertain, it will likely be strongly dependent on properties of the resident microbial communities and the local soil conditions. As such, it is important to understand the natural abiotic and biotic variation that occurs within permafrost environments in order to accurately inform models aimed at predicting responses to change across these regions.

While both environmental conditions and microbial community composition of permafrost-affected soils are known to be highly variable, the degree to which variation in community composition is linked to physicochemical conditions of the soil is not well understood ^10^. In many non-permafrost soils, microbial community composition is shaped by physicochemical conditions, including pH ^11-13^, nutrient content ^14,15^, and soil moisture ^16-18^. Given that permafrost can support active microbial communities ^19,20^, it is reasonable to assume that similar factors may be important in structuring the permafrost microbiome. Alternatively, microbial community composition in these environments may be decoupled from physicochemical conditions that are found to be important in non-permafrost soils, and may instead be similarly shaped by the shared constraints imposed by prolonged freezing. Understanding how microbial communities are shaped by environmental conditions represents an important knowledge gap in permafrost microbiology.

While resolving drivers of community composition in permafrost environments will improve fundamental understanding of the microbiology of these extreme ecosystems, there is also practical importance in resolving how pre-thaw conditions may be used as predictors of system level response to thaw. Earth system models that integrate aspects of microbial community composition and function are gaining support to improve understanding of terrestrial carbon cycling and predictions about the fate of soil carbon in response to environmental change ^21,22^. However, permafrost environments bring a high level of complexity that is difficult to generalize in current models, because soil type, soil conditions, and carbon composition may all have important impacts on post thaw dynamics and carbon transformation ^23-26^. Additionally, the composition of pre-thaw communities may be a strong determinant of post-thaw processes, as permafrost microbial communities are expected to respond rapidly to thaw ^27,28^, and the abundance of particular taxa and functional genes can be important predictors of process rates, such as methanogenesis ^26,29,30^ and iron reduction ^31^. These findings underscore the importance of integrating knowledge of the physical environment, the chemical nature of the organic matter pool, and the structure and function of permafrost microbial communities to accurately predict rates of carbon metabolism in these systems. Spatially explicit studies capturing measures of soil heterogeneity are, therefore, necessary to inform models aimed at predicting microbial community responses to permafrost thaw and carbon fate in these environments.

The purpose of this work was to resolve the factors and processes that govern microbial community structure in permafrost-affected soils. We hypothesize that the factors and processes shaping permafrost microbial communities differ from those shaping non-permafrost soil communities, and reflect the unique constraints of the permafrost environment. We characterized patterns of microbial community composition along landscape gradients in the boreal forest ecosystem of the Caribou Poker Creek Research Watershed (CPCRW) near Fairbanks, AK. We examined the influence of dispersal and selection on patterns of community composition and evaluated the importance of permafrost physicochemical conditions, including soil organic matter composition and thermodynamic properties, as deterministic factors. As the first landscape-scale survey linking permafrost community composition to environmental variability, this work provides mechanistic understanding of the controls on permafrost communities. This understanding can, in turn, inform models aimed at predicting permafrost microbial community characteristics and responses to thaw.

## Materials and Methods

### Sample collection and processing

Samples were collected along a hydrologic gradient in the Caribou Poker Creek Research Watershed (CPCRW). CPCRW is a long-term ecological research (LTER) site and is representative of the discontinuous permafrost regions of interior Alaska (http://www.lter.uaf.edu/research/study-sites-cpcrw). The sampling site was located on a gentle southeast-facing slope. To efficiently capture spatial variation at the landscape scale, we used a cyclic sampling design, as opposed to regular grid spacing ^32^. A 3/5 cyclic sampling design with 4 m grid cells was employed along four replicate transects for 104 m: starting at the lowest elevation, transects were sampled at 0, 4, 12, 20, 24, 32, 40, 44, 52, 60, 64, 72, 80, 84, 92, 100, and 104 m. Four replicate transects ran parallel to each other based on a 2/3 cyclic sampling design with 10 m grid cells for 40 m (Figure 1).

**Figure 1:**
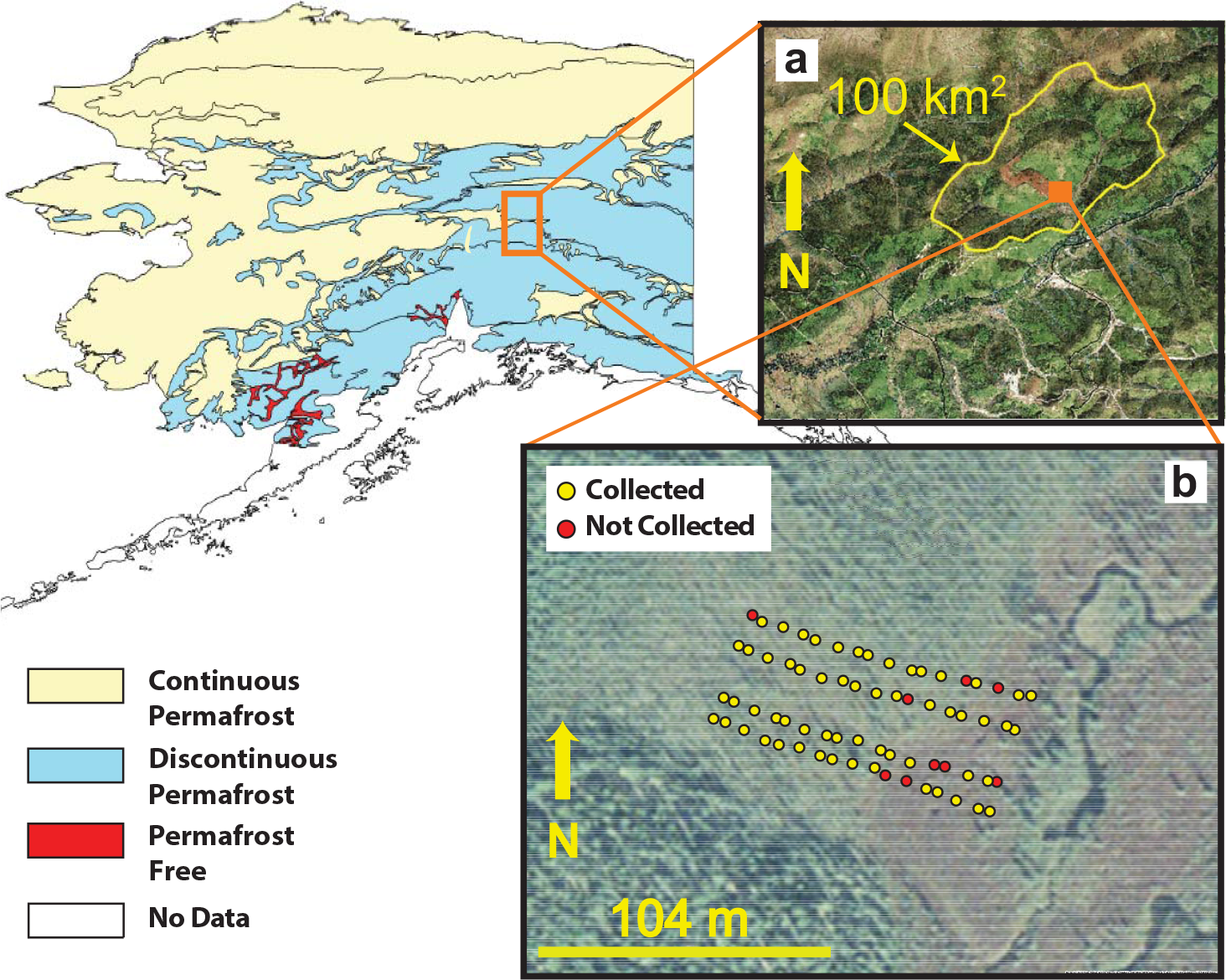
Map of Alaska indicating the location of a) the Caribou Poker Creek Research Watershed (CPCRW) and b) the location of the sample sites along each transect. Yellow dots indicate where samples were taken and included in the final analysis, while red dots indicate landscape positions where samples could not be recovered or where samples were compromised during processing, such that they were excluded from the final analysis.

At each sampling position, permafrost cores were collected using a SIPRE coring auger (John’s Machine Shop, Fairbanks, AK). Samples were wrapped in aluminum foil and packed in coolers on dry ice until they could be stored at −20 °C at the University of Alaska, Fairbanks, AK. Samples were shipped on dry ice to Pacific Northwest National Laboratory in Richland, WA, where they were stored at ‒20 °C until further processing.

The top 3-12 cm of each core was removed using an ethanol sterilized chisel and the next 4-10 cm section of each core was taken for analysis. The exterior of each core section was removed using sterile razor blades, starting from a pristine surface of the core. Decontaminated cores were crushed and homogenized while frozen using a sterile stainless steel soil press in a walk-in −20 °C freezer. Homogenized frozen samples were partitioned aseptically for downstream analyses.

Samples to be stored anaerobically were immediately purged with nitrogen gas in 60 ml serum bottles sealed with butyl rubber stoppers, and all partitioned material was stored at ‒20 °C until analysis. A total of 59 samples were included in the final analyses, as cores could not be retrieved from some sample locations or were compromised during sample processing (refer to Figure 1).

### Physicochemical analyses

Soil water content was determined by drying 1-10 g of sample at 105 °C, and measuring mass loss after 48 hours: sample masses were determined after 24 and 48 h to ensure samples reached a constant mass in consecutive measurements. Water content was determined from the average of five replicate measurements per sample.

Total carbon and nitrogen content was determined from 30 mg freeze-dried, ground, and <2 mm sieved samples. Samples were analyzed on an Elementar vario El cube (Elementar, Germany). Values were determined from the average of triplicate measurements for each sample.

Samples for metals and anion analyses were prepared from freeze-dried, ground, and <2 mm sieved samples. For metals analysis, 1 g of sample was extracted with 10 ml of 0.5 N HCl shaking at 200 rpm for 2 h at room temperature. Anion extractions were completed as above, with 1 g of soil in 5-10 ml of deionized water. Metals (Fe, Mn, Mg, Cu, P, S) and anion (Cl^−^, SO_4_^−^, and NO_3_-) analyses were completed as previously described (Zachara et al., 2009).

Iron(II) content was determined by ferrozine assay ^33^. In an anaerobic chamber (Coy Laboratory Products, Grass Valley, MI), 10 ml 0.5 N HCl was added to 1 g of permafrost sample, and the vial was sealed and vortexed. Samples were extracted for 1 h and filtered through a 0.22 pore-size polyethersulfone (PES) syringe filter. Extracts were diluted in 0.1 N HCl and 100 μl was added to 1 ml ferrozine; after 5 min, the absorbance at 562 nm was measured on a Shimadzu Biospec-1601 spectrophotometer. Iron(II) concentrations were determined from a six-point standard curve ranging from 0 to 45 μM Iron(II). Samples were dried at 60 °C and weighed to determine the Iron(II) content by dry weight.

Organic acids and sugars were quantified in the same water extracts prepared for anion analysis, using an Agilent 1100 series HPLC (Agilent, Palo Alto, CA) with a 300 × 7.8mm Aminex HPX-87H column (Bio-Rad, Hercules, CA), a 0.008 N H_2_SO_4_ mobile phase with a flow rate of 0.6 ml/min and variable wavelength detector (VWD) at 210 nm for organic acids and refraction index detector (RID) for sugars. Samples were filtered through a 0.22 μm pore-size PES syringe filter and acidified by adding 10 μl of 2.5 N H_2_SO_4_ per ml. Concentrations were determined by comparison to peak areas of standards.

Soil texture was determined by measuring the gravel (> 2mm), sand (64 μm- 2 mm), and mud (silt and clay) (<64 μm) fractions of each sample. Briefly, 20 g of soil was dried at 60 °C, and the total dry weight determined. Samples were dry sieved through a 2 mm sieve, and the mass of the >2 mm fraction was used to calculate the gravel fraction. The <2 mm fraction was wet sieved through a 64 sieve, the fraction retained was dried and used to calculate the sand fraction, while the <64 μ.m fraction was dried and used to calculate the mud fraction.

Soil pH was determined on a Denver Instrument model 215 pH meter (Denver Instruments, Bohemia, NY) by slurry of 1 g soil in 2 ml MilliQ water (Millipore Sigma, St. Louis, MO).

### Organic matter composition determination by FT-ICR-MS

Organic matter was extracted from bulk soil sequentially with water, methanol and chloroform as described previously ^34,35^. Briefly, organic matter extracts were prepared by adding 1 ml of solvent to 100 mg lyophilized and ground bulk soil and shaking for two hours on an Eppendorf Thermomixer in 2 mL capped glass vials. Samples were removed from the shaker and left to stand before centrifugation at 2000 rpm for 10 min and the supernatant was retained for analysis. The soil residue was dried with nitrogen gas to remove any residual solvent, and the extraction was repeated with each of the next two solvents. The chloroform and water extracts were diluted in methanol to improve electrospray ionization (ESI) efficiency and 20 μl was injected into the FTICR-MS. Samples were analyzed in triplicate for water extractions and chloroform extractions, and singly for methanol extractions.

A 12 Tesla Bruker SolariX FTICR-MS located at the Environmental Molecular Sciences Laboratory (EMSL) in Richland, WA, was used to collect high-resolution mass spectra of the organic matter in the extracts. A standard Bruker ESI source was used to generate negatively charged molecular ions. Samples were introduced directly to the ESI source at a flow rate of 3 μl/min. The ion accumulation time was varied, from 0.1 s to 0.5 s, to account for differences in C concentration between samples and to maintain a final dissolved organic carbon concentration of 20 ppm. The instrument was externally calibrated weekly with a tuning solution from Agilent (Santa Clara, CA), which calibrates to a mass accuracy of <0.1 ppm. Two hundred scans were averaged for each sample and internally calibrated using OM homologous series separated by 14 Da (-CH_2_ groups). The mass measurement accuracy was less than 1 ppm for singly charged ions across a broad m/z range (i.e. 200 <m/z <1200). To further reduce cumulative errors, all sample peak lists for the entire dataset were aligned to each other prior to formula assignment to eliminate possible mass shifts that would impact formula assignment. Putative chemical formulas were assigned using in-house software based on the Compound Identification Algorithm (CIA) ^36^, and modified as previously described ^37^. Chemical formulas were assigned based on the following criteria: S/N >7, and mass measurement error <1 ppm, taking into consideration the presence of C, H, O, N, S and P and excluding other elements. Peaks with large mass ratios (m/z values >500 Da) were assigned formulas through the detection of homologous series (CH_2_, O, H_2_). Additionally, to ensure consistent assignment of molecular formula the following rules were implemented: one phosphorus requires at least four oxygens in a formula and when multiple formula candidates were assigned the formula with the lowest error and with the lowest number of heteroatoms was picked.

For all analyses, peak intensities were converted to presence/absence and peaks observed in any of the triplicate measurements were included as present. Compound classes were assigned to chemical formulas based on molar O:C and H:C ratios, determined from analysis of van Krevelen diagrams. The Gibbs energies of the oxidation half reaction (ΔG°_Cox_) of each compound was derived based on the nominal oxidation state of carbon (NOSC) as previously described ^38^. The average ΔG°_Cox_ of the carbon pool was determined for each sample extraction: the median values were used for the methanol (ΔG°Cox(MeOH)) and chloroform (ΔG°_Cox_(CHC1_3_)) extracts due to highly skewed distributions, while the ΔG°_Cox_ was normally distributed for water extracts (ΔG°_Cox_(H_2_O)) such that the mean values were used.

### Microbial community analyses

Total community DNA was extracted from 0.25 g of each sample using the MoBio Power Soil DNA Isolation Kit (MoBio Laboratories, Carlsbad, CA), according to manufacturer’s instructions. Additional cleanup and concentration of DNA extracts was completed using the Zymo ZR-96 Genomic DNA Clean and Concentrator-5 kit (Zymo Research Corporation, Irvine, CA). PCR amplification of the V4 region of the 16S rRNA gene was performed as previously described ^39^, with the exception that the twelve-base barcode sequence was included in the forward primer. Amplicons were sequenced on an Illumina MiSeq using the 500 cycle Miseq Reagent Kit v2 (Illumina Inc., San Diego, CA), according to manufacturer’s instructions.

Raw sequence reads were demultiplexed using EA-Utils ^40^ not allowing any mismatches in the barcode sequence. Reads were quality filtered with BBDuk2 ^41^ to remove adapter sequences and PhiX with matching kmer length of 31 bp at a hamming distance of 1. Reads shorter than 51 bp were discarded. Reads were merged using USEARCH ^42^ with a minimum length threshold of 175 bp and maximum error rate of 1 %. Sequences were de-replicated and clustered using the distance-based, greedy clustering method of USEARCH at 97 % pairwise sequence identity among operational taxonomic unit (OTU) member sequences. Taxonomy was assigned to OTU sequences at a minimum identity cutoff fraction of 0.8 using the global alignment method implemented in USEARCH across RDP trainset version 15. OTU seed sequences were filtered against RDP classifier training database version 9 to identify chimeric OTUs using USEARCH. De novo prediction of chimeric reads occurred as reads were assigned to OTUs. OTU count tables were randomly resampled to 17899 sequences and OTUs that could not be assigned at the kingdom level were removed.

### Statistical analysis

The environmental variables consisted of all physicochemical variables and the average ΔG°_Cox_ for each FTICR extraction (mean for ΔG°_Cox_(H_2_O) and median for ΔG°_Cox_(MeOH) and ΔG°_Cox_(CHCh). Missing data were replaced by the geometric mean of values for a given variable, or the arithmetic mean in the case of the lactate data, which had numerous zero values. Data for water content, Cl, SO_4_, NO_3_, Fe(total), Mn, Mg, Cu, P, S, Fe(II), C, and N were log10(x) transformed, and data for lactate, formate, and acetate concentrations were log10(x+1) transformed. Data for pH, gravel, sand, mud, ΔG°_Cox_(H_2_O), ΔG°_Cox_(CHCh), and ΔG°_Cox_(MeOH) were not transformed.

Principal components analysis (PCA) was used to assess variation in environmental variables across the landscape using the *princomp* function in R ^43^. All variables were scaled by subtracting the mean and dividing by the standard deviation prior to analysis. Scores of all principal components (PCs) and variable loadings along each PC were extracted for downstream analyses. Variable loadings along each PC were used to assess the importance of individual variables to each PC.

Analyses of community diversity and composition were completed using the *‘vegan’* package ^44^ in R. Shannon diversity estimates were completed based on the resampled OTU counts. OTU abundances were Hellinger transformed prior to all other compositional analyses. Non-metric multidimensional scaling was used to examine the community variation between samples based on Bray Curtis dissimilarity, and environmental variables were fit as vectors in the final two-dimensional ordination to evaluate relationships between community and environmental variation.

Spatial analyses were completed in R. Kriging was used to interpolate and visualize spatial trends in both the environmental and biological data, using the autokrig function of the *‘automap’* package ^45^. Principal coordinates of neighbor matrices (PCNM) was used to create orthogonal spatial variables based on sample site locations ^46,47^. PCNMs were calculated as previously described ^48^. PCNM axes were used as explanatory variables in downstream analyses to examine the importance of spatial filters on community composition.

A redundancy analysis (RDA) model was used to relate community composition to environmental and spatial variation using the *vegan* package ^44^ in R. Due to collinearity between several environmental variables, PC scores extracted from the environmental PCA were used to represent environmental variables in the model. Forward stepwise model building based on adjusted R^2^ was carried out using all 23 PCs and all 15 positive PCNMs. The importance of each variable added to the model was assessed using variance partitioning based on RDA.

Null modeling was used to estimate the influence of ecological processes on community composition, as described previously ^49,50^. The influence of selection was estimated by evaluating the difference between the observed between-community mean-nearest-taxon distance (βMNTD) and the mean of the null distribution in units of standard deviation. Significant deviations from the null distribution were evaluated using the β–nearest taxon index (βNTI) and the signal for selection was expressed as the proportion of comparisons for which βNTI>2 or βNTI<-2, representing signals for variable selection and homogenous selection, respectively. Comparisons falling within the null distribution (2>βNTI>-2) represent compositional differences that do not arise from selection, and are instead attributable to dispersal limitation, homogenizing dispersal, or processes undominated by dispersal or selection. To assess the relative influence of these processes, a Raup-Crick metric incorporating species relative abundance (RC_bray_) was used to compare the observed and expected species turnover between communities. Significant deviations from the null distribution indicating greater than expected differences in community composition (2>βNTI>-2 and RC_bray_<0.95) were attributed to dispersal limitation, while those indicating less than expected differences in community composition (2>βNTI>-2 and RC_bray_¼-0.95) were attributed to homogenizing dispersal. Comparisons falling within the null distribution of both metrics (2>βNTI >-2 and 0.95>RC_bray_>-0.95) represent differences in community composition that were not strongly governed by selection or dispersal (i.e., the observed differences were ‘undominated’).

A regression modelling approach was used to identify the environmental variables that explain variation in the process estimates for total selection (variable and homogenous selection combined). Here, process estimates were generated for each community by finding the fraction of pairwise comparisons—between a given community and all other communities—falling into the process categories summarized above ^50^. Community-level estimates of total selection were then used as the dependent variable in an exhaustive model selection using Bayesian information criterion (BIC), performed in the *‘leaps’* package ^51^ in R.

Path analysis was used to estimate interactions among environmental variables predicted to influence total selection. A hypothetical model outlining expected relationships between variables was evaluated using the sem function of the *‘sem’* package ^52^ in R (Figure S1). Adjustments to the model were informed by modification indices, which suggest addition of paths to improve model fit, and were included based on logical evaluation of potential associations between variables.

### Code availability

Custom computer code used in the current study is available from the corresponding author on reasonable request.

### Data availability

Sequence data has been deposited in the European Nucleotide Archive (ENA), under accession number PRJEB23054 (http://www.ebi.ac.uk/ena/data/view/PRIEB23054). All other datasets generated and analyzed in the current study are available from the corresponding author on reasonable request.

## Results

### Environmental conditions and carbon composition

Permafrost characteristics were highly variable across the sampling area, and are summarized in Table S1. Samples ranged greatly in carbon content from 1.3 to 35.8 %, and nitrogen content co-varied strongly with carbon content (r^2^=0.98), ranging from 0.1 to 2.0 %. Soil texture was typically dominated by sand (average 63 %), but had substantial inputs of mud (average 35 %). All samples were mildly acidic, ranging from pH 4.9 to 6.7. Notably, permafrost samples across the site varied greatly in ice content, with gravimetric water content varying from 0.28 to 9.2 g(water)/g(dry soil). Fe(II) content, indicative of soil redox conditions, was also highly variable, spanning over two orders of magnitude from 0.07 to 12.9 mg/g(dry soil).

The compound classes assigned to FTICR peaks based on van Krevelen diagrams showed distinct peak profiles for each solvent extraction (Table S2). Water extractions recovered the highest percentage of compounds classified as lignin-, condensed hydrocarbon-, carbohydrate-, tannin-, and amino sugar- like compounds, while methanol and chloroform extractions recovered the highest percentage of compounds grouping to unsaturated hydrocarbon- and lipid- like compounds. Compounds grouping as peptide- or protein- like were recovered in all fractions, representing 6.61 %, 8.93 %, and 4.96 % in the water, methanol, and chloroform extracts, respectively. A large percentage of compounds in each extraction were not assigned to a compound class (25-48 %). The ΔG°_Cox_ estimates from the FTICR profiles were tightly linked to the overall variation in FTICR compound classes for each extraction (Figure S2). The ΔG°_Cox_ estimates were, therefore, used to represent organic carbon profiles in downstream analyses, as they capture variation in organic carbon composition as a biochemically meaningful continuous variable that can be interpreted mechanistically.

PCA using all physicochemical variables revealed environmental gradients both within and between transects (Figure 2). The first two PCs accounted for nearly 58 % of the environmental variance, with 41 % captured on PC1 and 17 % on PC2. The strongest loadings along PC1 were for C content (−0.31), N content (−0.31), water content (−0.28), S content (−0.28), acetate (−0.28), ΔG°_Cox_(H_2_O) (−0.27), and formate (−0.27), while the strongest loadings along PC2 were for Fe(II) content (-0.37), soil texture fractions of mud (−0.37) and sand (0.35), pH (0.33), and P content (−0.33).

**Figure 2:**
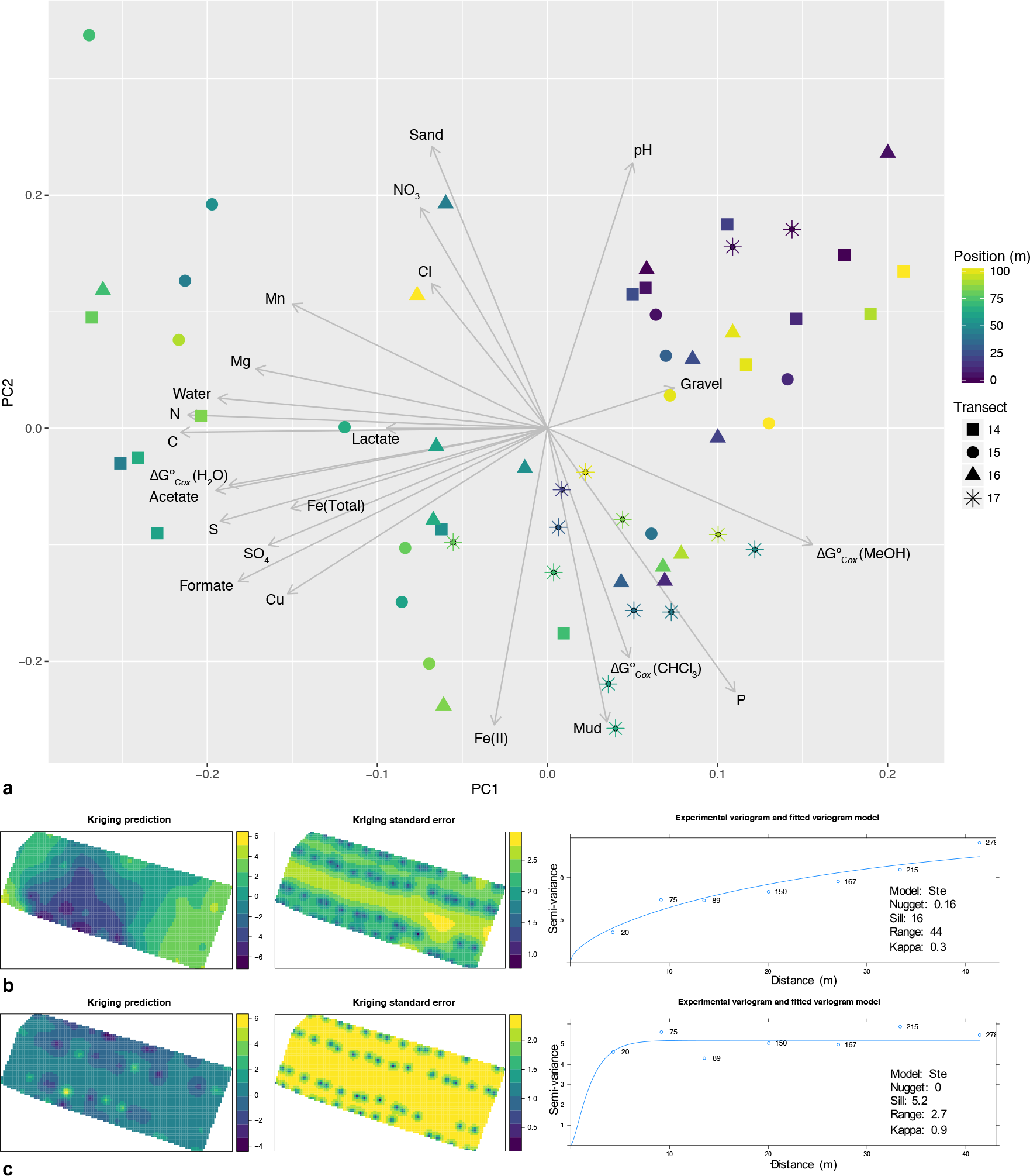
a) Principal components analysis (PCA) representing environmental variation between samples, and Kriging predictions of spatial patterns of environmental variation based on b) PC1 and c) PC2 scores across the sampling area (n=59).

### Microbial community composition

Bacterial sequences grouped to a total of 45 phyla or candidate phyla, and 11 phyla were represented at >1 % of the total community (Figure S3). Based on the average number of sequences in each sample grouping to bacterial phyla, communities were dominated by Proteobacteria (23.9 %) (particularly Beta- (10.9 %), Alpha-(5.9 %), Delta- (5.3 %), and Gamma- (1.5 %) proteobacteria), Acidobacteria (16.9 %), Verrucomicrobia (13.4 %), Actinobacteria (9.9 %), Chloroflexi (9.8 %), Bacteroidetes (8.5 %), Gemmatimonadetes (5.0 %), Planctomycetes (1.9 %), Nitrospirae (1.5 %), Parcubacteria (1.4 %), and Firmicutes (1.1 %). Bacterial sequences grouping to other phyla and bacterial sequences that could not be classified at the phylum level represented 4.8 % and 0.8 % of the total sequences, respectively.

Approximately 1.2 % of sequences were classified as Archaeal, with 79 % of these sequences grouping to the phylum Euryarchaeota. Sequences within the Euryarchaeota grouped predominantly within the Methanomicrobia and Methanobacteria.

### Patterns of community composition

Community composition showed non-random spatial structure (Figure 3), and was explained by both environmental variables (PCs) and, to a lesser extent, spatial variables (PCNMs). Stepwise model selection supported a model with fifteen variables, which fit the data with an adjusted R^2^=0.48; however, variance partitioning showed that many of these variables contributed only incrementally to improving model fit (Figure S4). A model incorporating the first two variables from the selected model (PC2, PC1) fit the data with an adjusted R^2^=0.34, and subsequent addition of the remaining variables retained in the selected model improved the adjusted R^2^ by 0.02 (PCNM4) or less (all other variables) (Figure 4). We therefore focused our interpretation on the model including PC2, PC1, and PCNM4. The variable loadings on the PCs selected in the model indicated several environmental factors were related to community composition (see Figure 2 for relationships between environmental factors along PC1 and PC2).

Univariate regression of factors with the strongest loadings along PC1 and PC2 showed that alpha diversity and the relative abundances of particular taxa were significantly associated with one or more of these environmental variables (Table S3).

**Figure 3:**
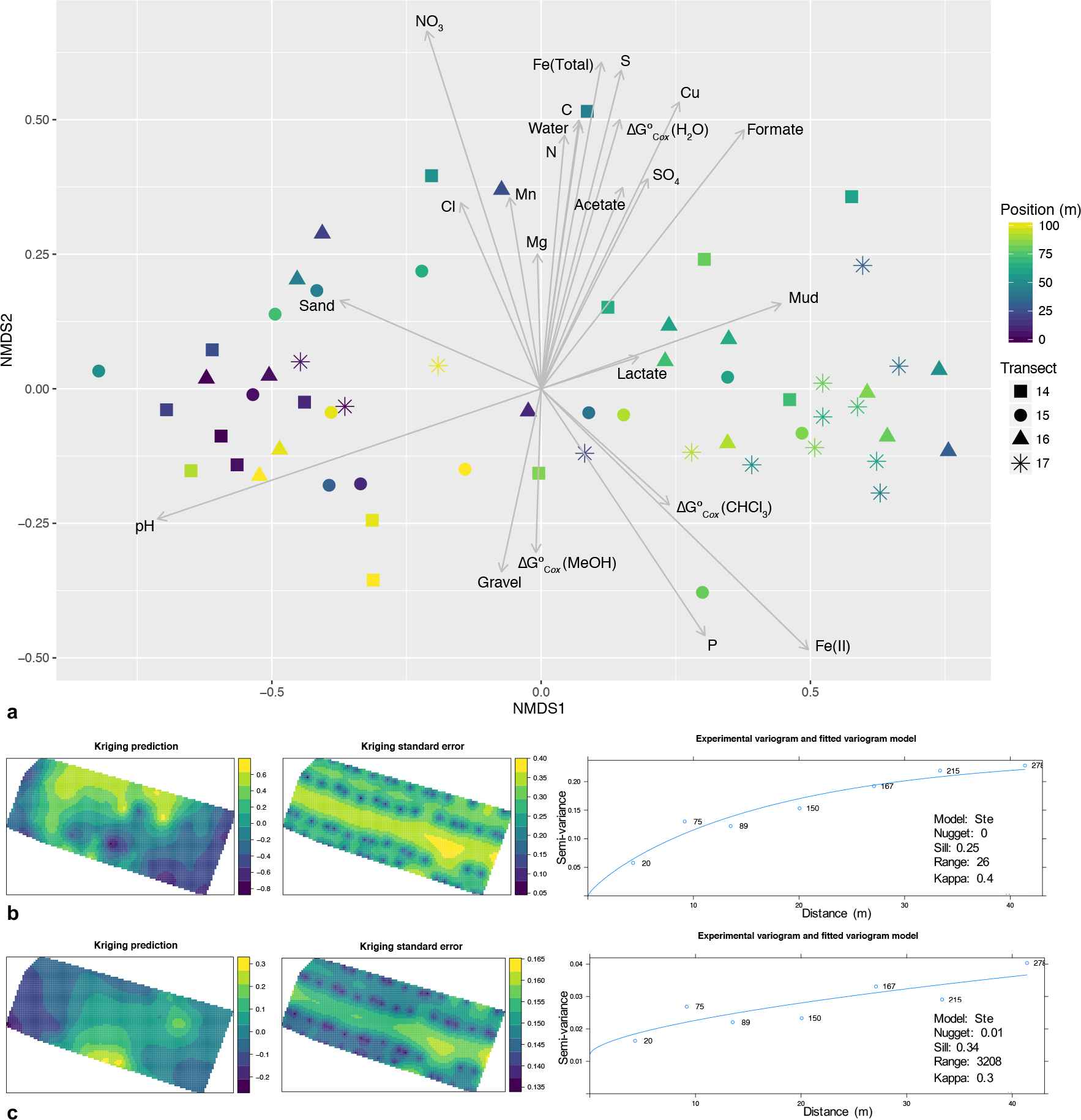
a) Non-metric multidimensional scaling (NMDS) plot representing the Bray Curtis dissimilarity in microbial community composition between samples, with environmental vectors overlaid, and Kriging predictions of spatial patterns of community composition based on b) NMDS1 and c) NMDS2 scores across the sampling area (n=59).

**Figure 4:**
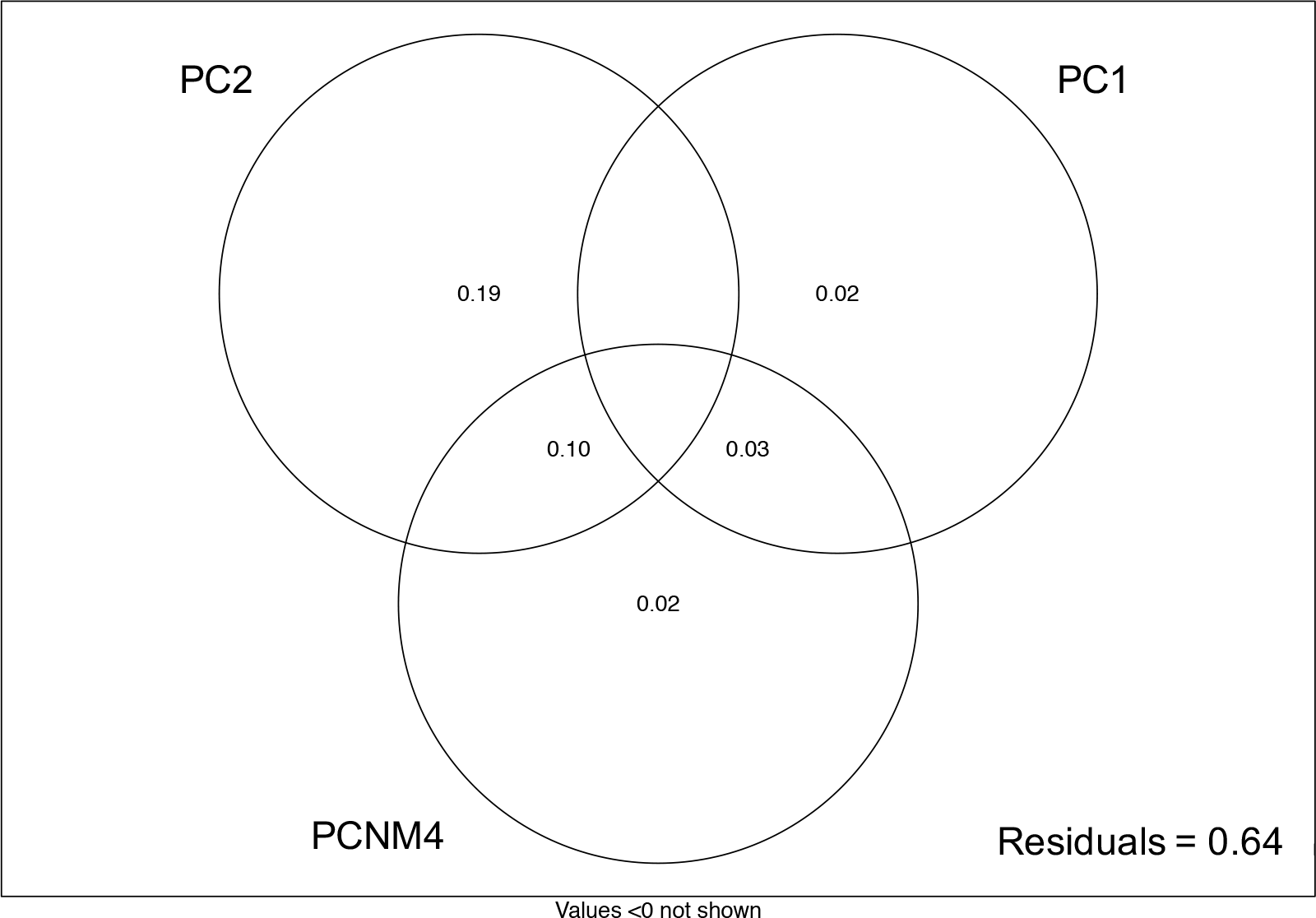
The proportion of variation in microbial community composition explained by the environmental and spatial variables selected in forward stepwise model building (n=59): including additional variables improved the adjusted R^2^ of the model by <0.02.

### Null model analyses

Null modeling revealed signals for variable selection (βNTI>2), homogenous selection (βNTI<-2), dispersal limitation (2>βNTI>-2 and RC_bray_>0.95), homogenizing dispersal (2> βNTI>-2 and RC_bray_<-0.95), and processes undominated by dispersal or selection (2> βNTI>-2 and 0.95> RC_bray_>-0.95) (Figure 5). Values from 0 to 1 indicating the relative influence of each process on the observed variation in community composition revealed the strongest signal was for dispersal limitation (0.36), and the lowest signal was for homogenizing dispersal (0.05). The signal for homogenous selection (0.21) was slightly higher than for variable selection (0.16), contributing to a signal of 0.37 for total selection. Variation not accounted for by dispersal or selection accounted for the remaining signal of 0.23.

**Figure 5:**
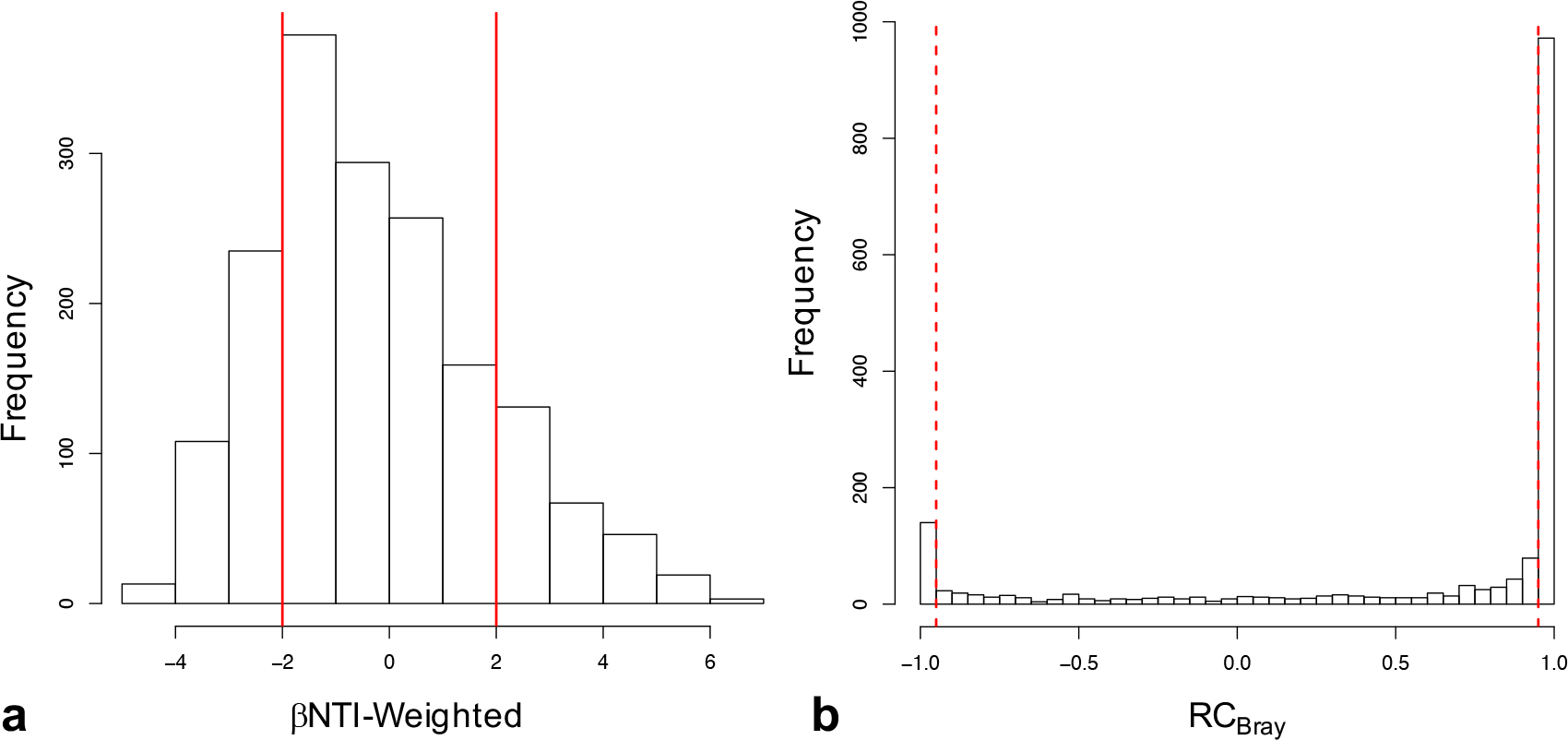
Histograms representing the observed distribution of comparisons based on a) βNTI and RC_bray_. Red lines represent the significance thresholds, whereby values outside their bounds are significantly different from the null distribution (n=59).

Regression model selection indicated Fe(II) was the most important environmental variable influencing variable selection, and Fe(II) was significantly associated with the relative influence of total selection by univariate regression (R^2^=0.14, p=0.003).

### Path analysis

Given the relationship observed between Fe(II) and total selection, we proposed a path model in which variables that reflect energetic constraints on microbial activity may influence total selection indirectly, through relationships mediated by Fe(II) content (See Figure S1). Our initial model was not consistent with the data (X^2^=37.2, d.f.=9 p=2.4×10^−5^), and was revised to better reflect relationships between the variables. All paths in the initial model were retained in the final model, and modification indices supported the addition of a path from soil carbon content to nitrate content. The final model did not differ significantly from the data (X^2^=10.3, d.f.=8, p=0.25) and explained 14.5 % of the variation in total selection and between 37 and 68 % of the variation in other endogenous variables (Figure 6). The direct effect of Fe(II) was the strongest total effect on total selection, while organic acid content had the strongest indirect effect on total selection (Table 1).

**Figure 6:**
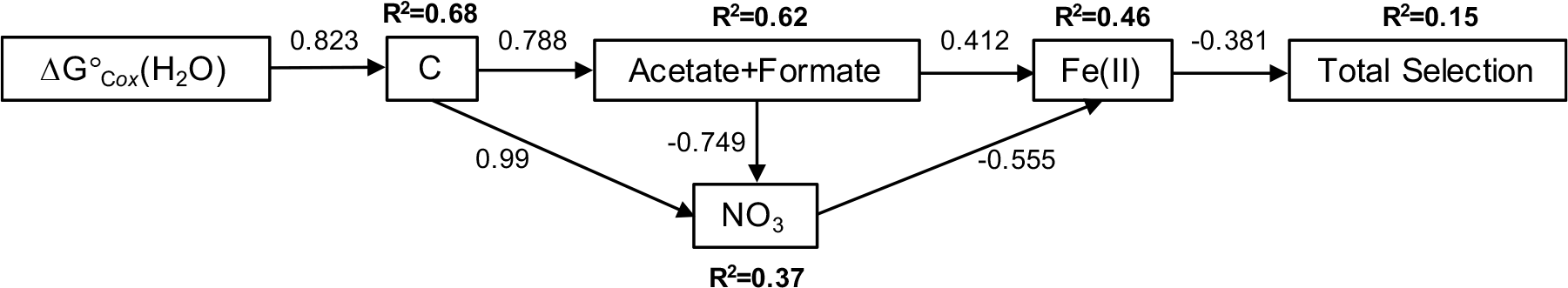
Final structural equation model (X^2^=10.3, d.f.=8, p=0.25) representing relationships between variables hypothesized to deterministically influence community composition (n=59). Values alongside arrows represent standardized path coefficients, and the variation explained for endogenous variables is indicated above each variable. All paths are significant.

**Table 1:** Standardized direct effects, indirect effects, and total effects of environmental factors on total selection.

|  | Direct Effects | Indirect Effects | Total Effects |
| --- | --- | --- | --- |
| $\Delta G^\circ_{Cox}(H_2O)$ | | -0.023 | -0.023 |
| Carbon |  | -0.039 | -0.039 |
| Acetate+Formate |  | -0.315 | -0.315 |
| Nitrate |  | 0.211 | 0.211 |
| Fe(II) | -0.381 |  | -0.381 |

## Discussion

Previous studies have demonstrated that permafrost soils contain diverse and varied communities that are likely active *in situ*; however, studies of permafrost microbiology typically suffer from low sample sizes, limiting the ability to examine ecological relationships that may influence community structure and function. To address this knowledge gap, we characterized patterns of environmental variation and microbial community composition in a boreal forest ecosystem across landscape gradients. By employing a well-replicated and geostatistically-informed sampling design, we have provided the first characterization of ecological processes driving landscape scale spatial structure of permafrost microbial community composition. Through this work, we show that patterns of both environmental characteristics and microbial community composition can be highly variable over short distances and exhibit non-random spatial structure, with patterns of community composition driven by deterministic and neutral processes that arise primarily from the physical constraints of the permafrost environment.

### Spatial structure of permafrost physicochemistry

The degree of heterogeneity in soil physicochemical and organic matter characteristics observed over the study area was striking, likely reflecting spatially structured variation in thaw history and organic matter deposition. We observed a non-linear spatial trend in environmental variation, with samples at the extreme ends of the transects found to be more similar to each other than to those in the middle of the transects, most notably in terms of water content, pH, carbon and nitrogen content, ΔG°_Cox_(H_2_O), and organic acid content (acetate and formate). Higher water content, which was observed predominantly through the middle of the transects, may represent ice inclusions in the transition zone near the surface of the permafrost, formed during more recent thaw events ^53^. Total carbon and nitrogen content, ΔG°_Cox_(H_2_O), and the abundance of organic acids were also highest through the middle of the transects, which may reflect more substantial deposition of undecomposed plant material from the active layer into the upper permafrost. The proportion of organic compounds grouping to lignins, carbohydrates, and amino sugars were highest in water extracts from the middle of the transects, and substantial deposits of fibric material were observed in many of these same samples. This undecomposed plant matter likely contributes high ΔG°_Cox_ compounds, such as lignin-like compounds, increasing the average ΔG°_Cox_ of the carbon pool.

We suggest that the higher organic acid concentrations observed in the middle of the spatial domain arise from the fermentation of labile organic compounds derived from deposited plant matter. If the most thermodynamically favorable compounds are preferentially fermented, this would further increase the average ΔG°_Cox_. In sediments, a net accumulation of organic acids is observed when fermentation rates exceed respiration rates ^54^, and acetate and C_1_ compounds are the dominant organic products of anaerobic metabolism in northern wetlands and bogs ^55,56^. These products of anaerobic metabolism may accumulate in permafrost through equivalent processes.

### Microbial community composition and environmental correlates

Community composition across our study site shared similarities with permafrost communities reported previously from across the Arctic, suggesting a core permafrost microbiome may be selected for by shared environmental constraints across disparate locations. We observed high representation of Acidobacteria and Proteobacteria, consistent with permafrost communities reported from Sweden ^30^, and parts of Alaska ^31^, and high representation of Chloroflexi, which has recently been reported in other Alaskan permafrost samples ^28,31^. Archaeal communities represented only a small percentage of the libraries and were dominated by taxa grouping to methanogens in the phylum Euryarchaeota, which is consistent with previous reports from across the Arctic ^28,30,31,57^. In contrast with previous studies, we saw high representation of Verrucomicrobia, which are globally abundant in soils ^58^, but have not been previously reported as dominant members of permafrost communities. Further comparisons of geographically distinct permafrost communities will require an increased number of studies employing well-replicated sampling designs and the adoption of standardized analytical techniques within the field ^7,10^.

Permafrost communities across the study site were influenced by similar drivers to non-permafrost soil communities, however several relationships were indicative of the unique constraints of the permafrost environment. Diversity was best described by a positive linear relationship with pH, which is consistent with trends observed in non-permafrost soils ^11-13^. The overall variation in community composition showed a clear relationship with environmental variation (PCs), although the particular environmental variables influencing community variation were not clear from the RDA model selection. Despite a relatively broad range of pH values across samples, no strong relationship between pH and community composition was observed. The relative abundance of the dominant phyla also varied significantly with numerous environmental variables, however these relationships were atypical of trends observed in surveys of non-permafrost soils. For example, at the phylum level, Acidobacteria are typically negatively associated with pH, while Bacteroidetes and Actinobacteria typically have positive relationships with pH ^11^; however, we observed the opposite trends for both Acidobacteria and Bacteroidetes and no trend for Actinobacteria with soil pH. Selective constraints of the permafrost environment may limit the phylogenetic breadth of these taxa, altering phylum level trends from those observed in other soils. Additionally, other deterministic factors, such as soil redox conditions and soil organic matter composition, which also showed strong univariate relationships with the relative abundance of particular taxa, may be more important drivers of community structure in permafrost-affected soils.

### Ecological processes influencing community composition

We employed a null modelling approach to evaluate the degree to which deterministic processes drive community variation and to resolve the variables most likely to be causally influencing composition. Patterns of community composition arise from a combination of deterministic and stochastic process ^59^ and the relative importance of these processes vary between systems. Null modeling provides a valuable tool to disentangle the influence of individual processes on patterns of microbial distribution ^50,60^. This approach has significant advantages over the RDA models, which cannot estimate relative contributions of assembly processes and did not reveal specific environmental variables that drive spatial variation in community composition.

Null modeling revealed a strong signal for dispersal limitation combined with a very weak signal for homogenizing dispersal, indicative of very restricted movement of microorganisms within the permafrost. The signal for dispersal limitation was stronger than for either homogenous or variable selection, and was effectively equivalent to the value for total selection. Dispersal limitation may be an especially important process in permafrost-affected soils, where microorganisms remain frozen in place for prolonged periods. Significant dispersal events may therefore be restricted to the limited movement that occurs through cryoturbation. These constraints likely limit community mixing over very short distances, which would lead to the strong signal of dispersal limitation observed in our null model analyses.

Given strong dispersal limitation, we expected that the spatial PCNM variables would explain significant variation in community composition, independent of environmental variation. This expectation was not met, however, with PCNM axes explaining little variation in the RDA model. The lack of a strong spatial signal in the RDA model indicates that the influence of spatial processes manifest below the spatial resolution of our sampling, consistent with very restricted movement of microorganisms through permafrost.

We found 37 % of the total community variation in permafrost community composition was explained by selective processes, and that soil characteristics associated with Fe(II) content are likely the most important environmental variables deterministically influencing community composition. Soil Fe(II) accumulates in anaerobic soils through the reduction of Fe(III), and iron reduction can contribute substantially to respiration in Arctic soils ^61,62^. A recent multi-omic analysis of Alaskan permafrost reported high representation of proteins annotated to iron-reducing taxa and the expression of genes annotated as cytochromes central to iron-reduction, suggesting iron-reducing taxa were likely active *in situ* ^31^. Importantly, Fe(III) reduction competes with other anaerobic processes, and suppresses less thermodynamically favorable methanogenic pathways ^63^. The relationship between Fe(II) and total selection indicates that soil redox conditions and thermodynamic constraints on microbial metabolism are likely to be the primary selection pressures that deterministically govern community composition.

These findings suggest that the stability of the permafrost environment strongly influences community structure and function, directly by restricting community mixing and indirectly by influencing the selective landscape, as electron donors and acceptors are depleted and infrequently replenished. This contrasts strongly with non-permafrost soils, in which communities are presumed to be well-dispersed through aeolian ^64^ and hydrologic process ^65^, nutrient fluxes are dynamic ^66,67^, and communities are thought to be shaped predominantly by selection ^68,69^.

Permafrost community structure and function, therefore, appear to be uniquely influenced by a balance between dispersal limitation imposed by frozen soil and deterministic selection arising primarily from thermodynamic constraints.

### Thermodynamic constraints

The thermodynamic constraints driving selection arise from the composition of both the organic matter and terminal electron acceptor pools, as outlined in our final SEM. We suggest that total carbon content accrues in the form of less favorable organic matter, as stocks of more favorable organic compounds are depleted; in turn, a relationship emerges wherein soil carbon content increases with ΔG°o« (higher values indicate lower favorability ^38^). Further, organic acids are expected to accumulate in these soils through anaerobic metabolism, as labile carbon is fermented. We suggest that these organic acids support nitrate and Fe(III) reduction, such that organic acid content is negatively associated with nitrate, and positively associated with Fe(II). Additionally, a negative relationship between nitrate and Fe(II) likely arises because Fe(III) reduction is less energetically favorable than nitrate reduction. Modification indices supported a positive association between total soil carbon and nitrate content in the final model: the positive association between total carbon and nitrate is consistent with our interpretation of higher carbon content resulting from accumulation of organic molecules that are less thermodynamically favorable for microbially-driven organic carbon oxidation. In this case, higher total carbon reflects less favorable carbon, which would result in lower rates of nitrate reduction that depend on the oxidation of organic carbon, and thus a positive carbon-nitrate relationship. We note that such inferences should be interpreted as speculative, given that controlled experiments were not conducted.

### Conclusions

Our findings support the hypothesis that permafrost microbial communities are shaped by factors that are distinct from those governing non-permafrost soil communities. We found that microbial distributions in permafrost are driven primarily by dispersal limitation imposed by frozen soil and deterministic selection arising from thermodynamic constraints of the permafrost environment. This contrasts sharply with non-permafrost soil communities, which are driven primarily by soil pH ^11-13^. These findings underscore the need for different mechanistic models predicting microbial community characteristics in permafrost and non-permafrost soils, given the different processes governing these systems. Our findings suggest that predictive models of permafrost community composition will need to account for organic carbon thermodynamics, organic acid concentrations, and redox conditions, which may be informed by knowledge of landscape history. However, efforts to accurately predict community composition at the landscape-scale based solely on environmental characteristics may be limited due to the strong influence of dispersal limitation.

Our findings additionally suggest that changes in permafrost microbial community structure and function are likely to be drastic in response to thaw, as hydrologic changes mobilize organisms and nutrients, thereby relieving the primary constraints on communities. Community responses to change are also likely to be highly varied across landscapes, given the environmental and microbiological heterogeneity of permafrost-affected soils. Identifying how pre-thaw environmental and community characteristics influence post-thaw responses will be essential for accurately predicting ecosystem level responses to environmental change.

## Acknowledgments

This research was supported by the Microbiomes in Transition Initiative at Pacific Northwest National Lab (PNNL), a multiprogram national laboratory operated by Battelle for the U.S. Department of Energy, and was conducted under the Laboratory Directed Research and Development Program at PNNL. A portion of this research was conducted using PNNL Institutional Computing resources and at the Environmental Molecular Sciences Laboratory (EMSL), a national scientific user facility in Richland, WA. We thank Caroline Anderson and Alex Crump for their assistance with sample collection and field work. We thank Tom Wietsma for conducting carbon and nitrogen analyses and Tom Resch for conducting metals and anion analyses.

## Author Contributions

This work was conceived by EMB, JKJ, and JCS. EMB, DWK, EBR, and SJF completed sample processing and laboratory analyses. Processing of amplicon sequencing data was completed by JMB. MMT and RKC performed FTICR-MS analyses and data processing. Statistical analyses were completed by EMB, JCS, and LMB. EMB wrote the manuscript, with input from all co-authors.

